# RNA editing of AZIN1 coding sites is catalyzed by ADAR1 p150 after splicing

**DOI:** 10.1101/2023.04.17.537126

**Authors:** Yanfang Xing, Taisuke Nakahama, Yuke Wu, Maal Inoue, Jung In Kim, Hiroyuki Todo, Toshiharu Shibuya, Yuki Kato, Yukio Kawahara

## Abstract

Adenosine-to-inosine RNA editing is catalyzed by nuclear ADAR1 p110 and ADAR2, and cytoplasmic ADAR1 p150 in mammals, all of which recognize double-stranded RNAs (dsRNAs) as targets. Although its frequency is quite rare, RNA editing occurs in coding regions, which alters protein functions by exchanging amino acid sequences, and is therefore physiologically significant. In general, such coding sites are edited by ADAR1 p110 and ADAR2 prior to splicing, given that the corresponding exon forms a dsRNA structure with an adjacent intron. We previously found that RNA editing at two coding sites of antizyme inhibitor 1 (AZIN1) is sustained in *Adar1 p110/Aadr2* double knockout mice. However, the molecular mechanisms underlying RNA editing of AZIN1 remain unknown. Here, we showed that Azin1 editing levels were increased upon type I interferon treatment, which activated Adar1 p150 transcription, in mouse Raw264.7 cells. Azin1 RNA editing was observed in mature mRNA but not precursor mRNA. Furthermore, we revealed that the two coding sites were editable only by ADAR1 p150 in both mouse Raw264.7 and human HEK293T cells. This unique editing was achieved by forming a dsRNA structure with a downstream exon after splicing and the intervening intron suppressed RNA editing. Therefore, deletion of a nuclear export signal from ADAR1 p150, shifting its localization to the nucleus, decreased Azin1 editing levels. Finally, we demonstrated that Azin1 RNA editing was completely absent in *Adar1 p150* knockout mice. Thus, these findings indicate that RNA editing of AZIN1 coding sites is exceptionally catalyzed by ADAR1 p150 after splicing.

## Introduction

Post-transcriptional modifications increase protein diversity from the limited information encoded in the genome. Adenosine-to-inosine (A-to-I) RNA editing, which occurs in double-stranded RNA (dsRNA) structures, is such a modification with an estimate of more than 100 million sites in human transcripts (1–3). Adenosine deaminase acting on RNA 1 (ADAR1) and ADAR2 that contain dsRNA-binding and deaminase domains are active enzymes responsible for A-to-I RNA editing in mammals (4–7). Furthermore, ADAR1 is expressed as two isoforms: ADAR1 p150 that is driven by a type I interferon (IFN)-inducible promoter and possesses a Z-DNA/RNA-binding domain (Zα) and a nuclear export signal (NES) in the N-terminus; and N-terminally truncated ADAR1 p110 regulated by a constitutive promoter (8–13). Therefore, ADAR1 p150 is mainly localized in the cytoplasm under normal conditions and is abundant in lymphoid organs such as the thymus and spleen, whereas ADAR1 p110 and ADAR2 are localized in the nucleus and are abundant in the brain where ADAR1 p150 is expressed at its lowest level (6,14–18).

*Adar1* knockout (KO) mice (*Adar1^-/-^* mice), *Adar1 p150*–specific KO (*Adar1 p150^-/-^*) mice, and *Adar1* knock-in (KI) mice harboring an editing-inactive E861A point mutation (*Adar1^E861A/E861A^* mice) all show embryonic lethality with increased expression of IFN-stimulated genes (ISGs) (19–22). Such abnormal phenotypes are rescued by concurrent deletion of *Ifih1*-encoded melanoma differentiation-associated protein 5 (MDA5) or its downstrem mitochondrial antiviral signaling protein (MAVS) (21–24). In contrast, *Adar1 p110*–specific KO (*ADAR1 p110^-/-^*) mice show a high mortality rate during their early post-natal days without overexpression of ISGs, which are caused by RNA editing–independent functions (14). In addition, *Adar2* KO (*Adar2^-/-^*) mice die within 3 weeks after birth due to progressive seizures (25,26). Such lines of evidence indicate that cytoplasmic ADAR1 p150–mediated RNA editing prevents MDA5-sensing of endogenous dsRNAs formed within mRNA, leading to the overexpression of ISGs, by altering dsRNA structure, which differs from the roles of nuclear ADAR1 p110 and ADAR2. Of note, the expression of ADAR1 p150 is enhanced as an ISG under certain conditions, such as a viral infection (27).

Most RNA editing sites are present in inverted repetitive sequences, such as short interspersed elements (SINEs), which form long dsRNA structures, especially in 3’ untranslated regions (UTR) and introns (28–32). RNA editing at some of these sites is most likely critical for preventing MDA5 activation. In contrast, although the number of sites is extremely limited, RNA editing occurs in certain coding regions, which is termed “recoding” (33,34). In these cases, RNA editing can change amino acid sequences and alter the physiological properties of the resultant proteins, given that inosine is interpreted as guanosine by the translational machinery (35). For instance, RNA editing at the ADAR2-specific Q/R site of GluA2, a glutamate receptor subunit, substitutes glutamine (Q) for arginine (R) at this site, affecting the Ca^2+^ permeability of the receptor (36). Progressive seizures observed in *Adar2^-/-^* mice are attributed to the increased Ca^2+^ permeability caused by the loss of RNA editing at the Q/R site of GluA2 (25). RNA editing at such coding sites requires a dsRNA structure, which is usually formed within a single exon or between the editing site–containing exon and the editing (or exon) complementary sequence (ECS) located in an adjacent intron (37–40). As examples of the former, Kv1.1 I/V and GABRA3 I/M sites, both of which are ADAR2-preferential sites, are known (6,38,39). In contrast, RNA editing of BLCAP coding sites requires an ECS in the upstream intron, while the Q/R site of GluA2, the E/G site of CAPS1, and five sites in the serotonin 5-HT_2C_ receptor need an ECS in the downstream intron for their RNA editing (34,41–44). This type of RNA editing is catalyzed by ADAR1 p110 and ADAR2 in the nucleus before splicing.

We recently investigated ADARs responsible for RNA editing *in vivo* at all coding sites that are conserved between humans and mice (6). Most coding sites show higher editing in the brain where ADAR1 p110 and ADAR2 are abundant. However, E/E and S/G sites of antizyme inhibitor 1 (AZIN1) show higher editing in the spleen where ADAR1 p150 is relatively abundant. Furthermore, we found that the editing levels at these two sites of Azin1 mRNA are not reduced in *Adar1 p110^-/-^/Adar2^-/-^* mice (14), which further suggests that ADAR1 p150 is responsible for Azin1 RNA editing *in vivo*. However, the molecular mechanisms underlying this unique editing pattern of coding sites for AZIN1 protein remain unknown.

In this study, we found that RNA editing of Azin1 was enhanced upon type I IFN treatment *in vitro* and these sites were editable only by ADAR1 p150 in both human and mouse cell lines. Of note, we found that RNA editing of AZIN1 coding sites required a dsRNA structure formed between the editing site–containing exon and the downstream exon, and that the intron intervening between these two exons acted as a potent suppressor for RNA editing. Accordingly, RNA editing at Azin1 coding sites was absent in *Adar1 p150^-/-^* mice, which indicates that ADAR1 p150 is responsible for Azin1 RNA editing *in vivo*. These findings collectively indicate that AZIN1 is the first case in which coding sites are editable after splicing, modulating the degree of recoding via regulation of ADAR1 p150 expression.

## Results

### RNA editing of Azin1 coding sites is responsive for the treatment of type I interferon

To examine the molecular mechanisms underlying Azin1 RNA editing, we first investigated whether two editing sites, which are located in exon 11 (Ex11) of the *Azin1* gene (**Fig. 1A**), were responsible for type I IFN treatment *in vitro*. For this purpose, we stimulated mouse macrophage-like Raw 264.7 cells, which express both ADAR1 p110 and p150 at a comparable level, with IFNβ1 (**Fig. 1B**). As expected, the expression of ADAR1 p150, which is driven by the type I IFN–inducible promoter, was significantly increased whereas the expression of ADAR1 p110 was unaffected (**Fig. 1B–D**). This treatment did not affect the expression of Azin1 mRNA and AZIN1 protein (**Fig. 1E, F**). We subsequently examined editing levels at two editing sites of mature Azin1 mRNA. Although the editing ratios at E/E and S/G sites were ∼12% and ∼33%, respectively, under normal conditions, the editing levels of these two sites significantly increased to ∼22% and ∼51%, respectively, after type I IFN treatment (**Fig. 1G**). In contrast, no substantial RNA editing was detected in both sites in precursor mRNA that contained downstream intron 11 (In11) (**Fig. 1H**). These results suggest that RNA editing at Azin1 coding sites is IFN-responsive, depending upon the amount of ADAR1 p150, and these sites are edited after mRNA maturation.

**Figure 1.**
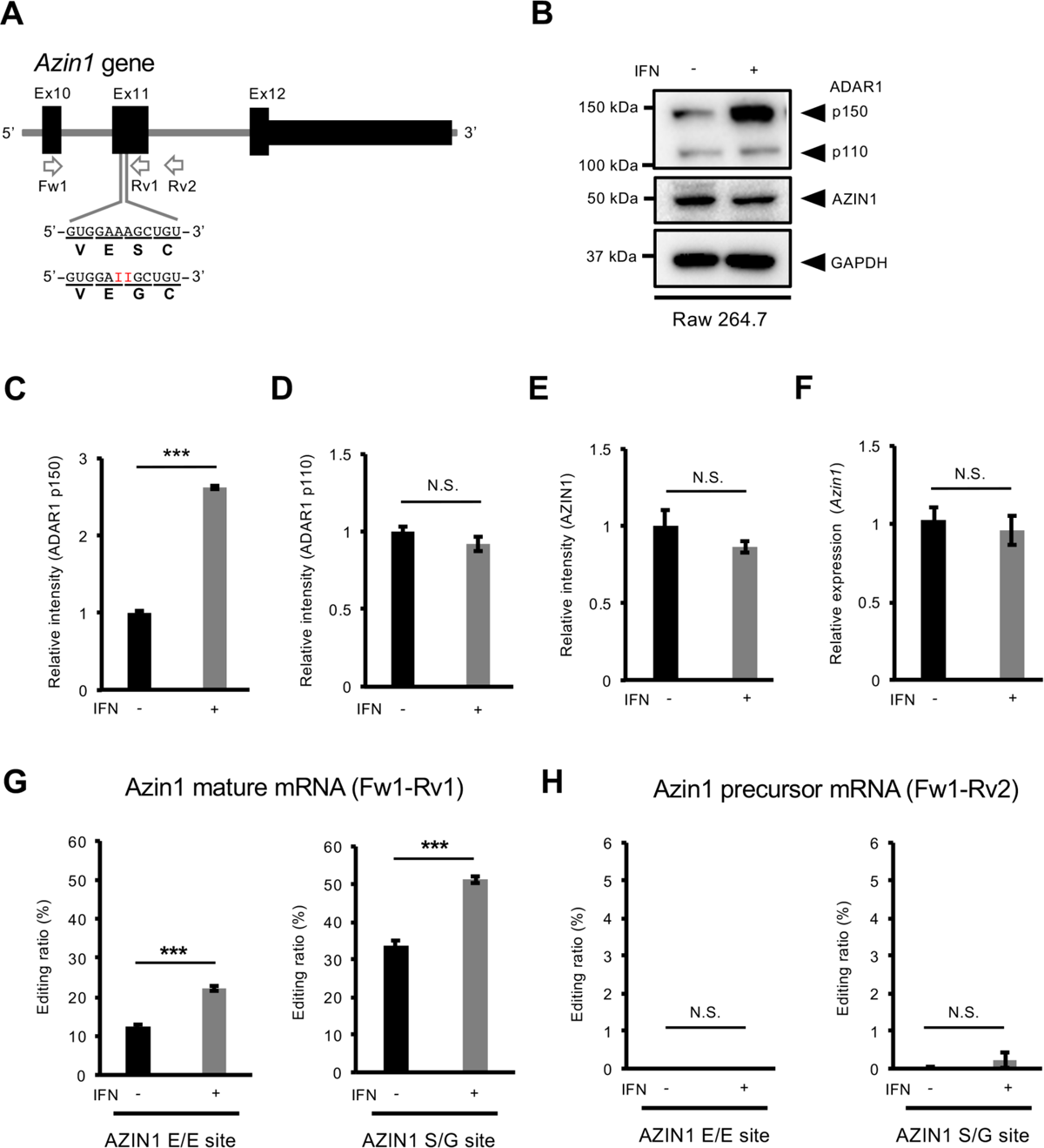
Induction of Azin1 RNA editing by IFN treatment. (**A**) Schematic diagram of the mouse *Azin1* gene, which is conserved with the human *AZIN1* gene. Two editing sites (shown as red Is) located in exon 11 (Ex11) and the resultant amino acid sequences are indicated below. The position of PCR primers is also shown by arrows. (**B**) The expression of ADAR1 p110, p150, and AZIN1 proteins in Raw 264.7 cells in the absence or presence of interferon (IFN) β1 treatment is shown. The expression of GAPDH protein is shown as a reference. (**C–E**) The band intensity of ADAR1 p150 (**C**), ADAR1 p110 (**D**), and AZIN1 (**E**) proteins in the absence or presence of IFNβ1 treatment was normalized to that of GAPDH and displayed as the mean ± SEM (n = 3 for each group; Student’s *t*-test, \*\*\**p* < 0.005, N.S., not significant) by setting the mean value of the intensities without IFNβ1 treatment as 1. (**F**) The relative expression of Azin1 mRNA in Raw 264.7 cells in the absence or presence of IFNβ1 treatment is shown. Values represent relative gene expression normalized to GAPDH mRNA and are displayed as the mean ± SEM (n = 3 for each group; Student’s *t*-test, N.S., not significant) by setting the mean value of expression without IFNβ1 treatment as 1. (**G, H**) RNA was extracted from harvested cells and subjected to reverse transcription (RT) followed by PCR using Fw1-Rv1 (**G**) or Fw1-Rv2 (**H**) rimer sets. The PCR products derived from mature (**G**) and precursor (**H**) Azin1 mRNA were used for the quantification of editing ratios at two sites of Azin1 mRNA in the absence or resence of IFNβ1 treatment. Values are displayed as the mean ± SEM (n = 3 for each group; Student’s *t*-test, \*\*\**p* < 0.005, N.S., not significant).

### Azin1 coding sites are editable by ADAR1 p150 but not ADAR1 p110 and ADAR2 *in vitro*

Next, we transfected a plasmid containing enhanced green fluorescent protein (EGFP)-tagged mouse ADAR1 (mADAR1) p110, p150 or mADAR2 into *Adar1/Adar2* double-knockout (A1/A2 dKO) Raw 264.7 cells, which were previously established (18). As previously reported, we observed that ADAR1 p110 and ADAR2 were predominantly localized in the nucleolus, while ADAR1 p150 was mainly detected in the cytoplasm (15,18,45–47) (**Fig. 2A**). After each ADAR was induced in A1/A2 dKO Raw 264.7 cells, we showed that each EGFP-tagged mADAR protein was expressed at the expected size (**Fig. S1A**) and then sorted GFP-positive cells by adjusting the expression of each mADAR protein using the intensity of GFP fluorescence. After RNA extraction from the sorted cells, editing levels were examined. We found that the ADAR2-specific S/G site of Nova1 was editable by only ADAR2 in this system (6) (**Fig. S1B**). In addition, Y/C and Q/R sites of Blcap mRNA, known ADAR1-target sites, were preferentially edited by ADAR1 p110 in addition to ADAR2, which could edit these sites to some extent (**Fig. S1C**). Given that RNA editing of Blcap coding sites requires ECS in the upstream intron (6,14,34), ADAR1 p150, which is mainly detected in the cytoplasm, had no ability to edit Blcap coding sites, as expected. In contrast, two sites of Azin1 mRNA were editable by ADAR1 p150 but not ADAR1 p110 and ADAR2 (**Fig. 2B**). This ADAR1 p150–mediated RNA editing was not detected in precursor Azin1 mRNA containing In11 (**Fig. 2C**).

**Figure 2.**
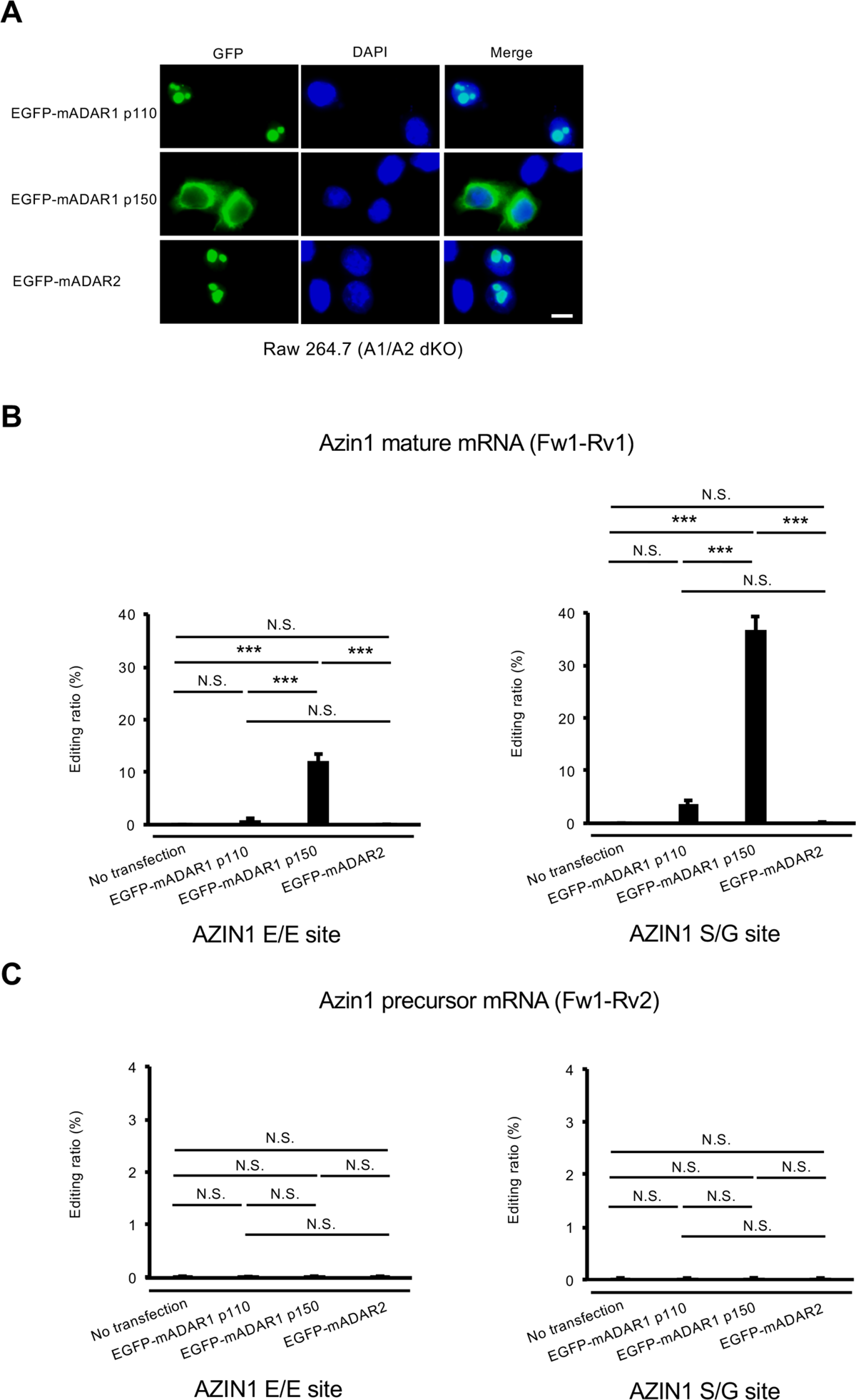
ADAR1 p150–specific RNA editing of mouse Azin1 mRNA. (**A**) The intracellular localization of EGFP-tagged mouse ADAR1 p110 (mADAR1 p110), mADAR1 p150, and mADAR2 in *Adar1*/*Adar2* double-knockout (A1/A2 dKO) Raw 264.7 cells was visualized by GFP fluorescence. Nuclei were stained with DAPI. Merged fluorescent images are shown in the right panels. Scale bar, 10 μm. (**B, C**) After the indicated EGFP-tagged ADAR isoforms were expressed in A1/A2 dKO Raw 264.7 cells, RNA was extracted and subjected to reverse transcription (RT) followed by PCR using Fw1-Rv1 (**B**) or Fw1-Rv2 (**C**) primer sets shown in Figure 1A. The PCR products derived from mature (**B**) and precursor (**C**) Azin1 mRNA were used for the quantification of editing ratios at two sites of Azin1 mRNA. Values represent the mean ± SEM (n = 3 for each group; Tukey’s honest significant difference test, \*\*\**p* < 0.005, N.S., not significant).

To examine the conservation of AZIN1 RNA editing between humans and mice, we transfected a plasmid containing HaloTag-fused human ADAR1 (hADAR1) p110, p150 or hADAR2 into human HEK293T cells. Of note, the editing level of AZIN1 coding sites was less than 2% under normal conditions, which most likely reflects the low expression of ADAR1 p150 in HEK293T cells in which ADAR1 p110 is relatively abundant (**Fig. S2A, B**). In these cells, the editing ratios at E/E and S/G sites of AZIN1 mRNA were increased to ∼15% and ∼30%, respectively, when ADAR1 p150 was forcibly expressed. In contrast, such an increase in RNA editing was not observed when ADAR1 p110 or ADAR2 was overexpressed (**Fig. S2B**). These results indicate that AZIN1 coding sites are editable *in vitro* only by ADAR1 p150 at the mature mRNA stage, which is conserved between human and mouse cell lines.

### dsRNA structure required for Azin1 RNA editing is formed with the downstream exon

ADARs recognize a dsRNA structure as a target. The editing sites in coding regions are usually located in a dsRNA structure formed within a single exon or between the editing site–containing exon and ECS in an adjacent intron (38-40,42-44). To identify the dsRNA structure required for Azin1 RNA editing, we created three mCherry-fused reporter constructs that contained the editing site–containing Ex11 and an upstream Ex10 and In10 (mEx10-In10-Ex11), Ex11 alone (mEx11), and Ex11 and a downstream In11 and partial Ex12 (mEx11-In11-Ex12), respectively (**Fig. 3A**). After each mCherry-fused reporter mRNA was expressed in Raw 264.7 cells, we investigated whether RNA editing occurred at the two sites in Ex11. No RNA editing was detected in both In10-spliced transcripts derived from an mEx10-In10-Ex11 reporter construct and transcripts derived from an mEx11 reporter construct (**Fig. 3B**). In contrast, we observed that E/E and S/G sites were edited at levels of 1.5% and 6.4%, respectively, under normal conditions, which increased to 4.2% and 16.5%, respectively, in the presence of type I IFN in In11-spliced transcripts derived from an mEx11-In11-Ex12 reporter construct (**Fig. 3B**). Although In11 was efficiently spliced out, we further examined RNA editing in In11-containing transcripts, which were derived from both precursor mRNA and mature mRNA with In11 retained, resulting in no detection of substantial RNA editing (**Fig. 3C, D**). These data suggest that a dsRNA structure is formed between Ex11 and Ex12 after the splicing of In11, which inhibits RNA editing of AZIN1 coding sites.

**Figure 3.**
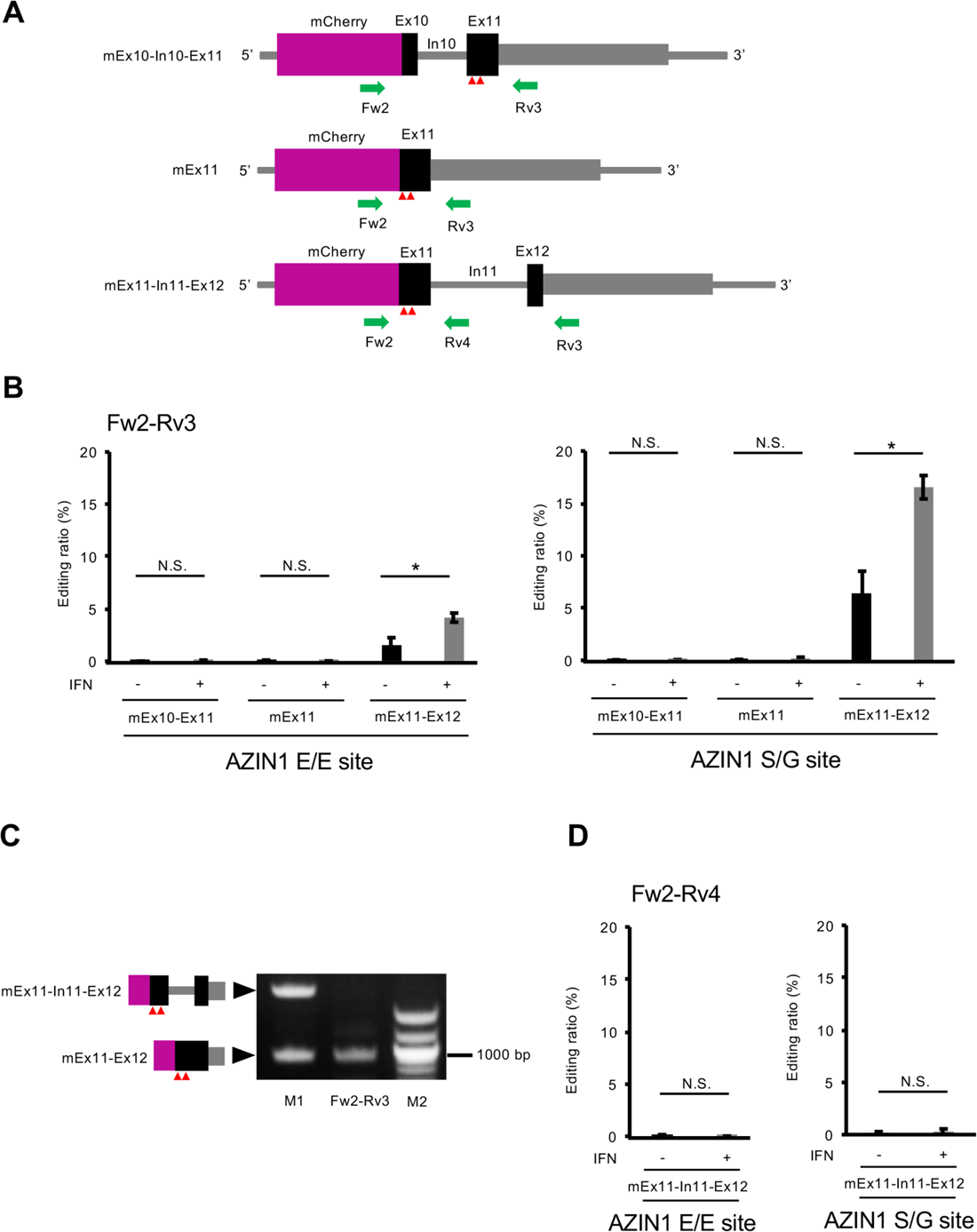
ADAR1 p150–mediated RNA editing of Azin1 after splicing. (**A**) Schematic diagrams of the reporter construct of the mouse *Azin1* gene encompassing exon 10 (Ex10), intron 10 (In10), and Ex11 (mEx10-In10-Ex11), that of Ex11 alone (mEx11), and that of Ex11, In11, and partial Ex12 containing only the coding region (mEx11-In11-Ex12), which is fused to the *mCherry* gene. The position of PCR primers is shown below. Two editing sites are indicated with red arrowheads. (**B**) After each mCherry-fused Azin1 reporter vector was transfected into Raw 264.7 cells, these were treated with interferon (IFN) β1. Then, RNA was extracted and subjected to reverse transcription (RT) followed by PCR using Fw2-Rv3 primer sets. The PCR roducts derived from mature Azin1 reporter mRNA were used for the quantification of editing ratios at AZIN1 E/E and S/G sites in the absence or presence of IFNβ1. Values represent the mean ± SEM (n = 3 for each group; Student’s *t*-test, \**p* < 0.05, N.S., not significant). (**C**) After an mCherry-fused mEx11-In11-Ex12 reporter vector was transfected into Raw 264.7 cells, RNA was extracted from the harvested cells and subjected to RT followed by PCR using Fw2-Rv3 primer sets. The resultant PCR products were subjected to electrophoresis with DNA size markers (M2, right lane) and position markers (M1, left lane). The expected sizes of the PCR roduct containing Ex12, In11, and partial Ex12, and that of Ex11, and partial Ex12 are 1,921 and 918 bp, respectively. (**D**) After an mCherry-fused mEx11-In11-Ex12 reporter vector was transfected into Raw 264.7 cells, cells were treated with IFNβ1. Then, RNA was extracted and subjected to RT followed by PCR using Fw2-Rv4 primer sets. The PCR products derived from recursor Azin1 reporter mRNA were used for the quantification of editing ratios at AZIN1 E/E and S/G sites in the absence or presence of IFNβ1. Values represent the mean ± SEM (n = 3 for each group; Student’s *t*-test, N.S., not significant).

Considering RNA editing of AZIN1 coding sites is conserved between humans and mice, the dsRNA structure required for this RNA editing is also likely conserved. In this regard, the conservation of In11 is relatively low, whereas the predicted dsRNA structure formed between Ex11 and Ex12 is highly conserved between humans and mice (**Fig. S3A, B**). To validate the dsRNA structure, we introduced four sequential point mutations into the predicted complementary region facing the editing sites in Ex12 of an mCherry-fused mEx11-In11-Ex12 reporter construct and expressed this mutant construct in Raw 264.7 cells (**Fig. 4A**). As expected, no substantial RNA editing was observed at E/E and S/G sites, regardless of IFNβ1 treatment (**Fig. 4B**). Taken together, these results indicate that the dsRNA structure formed between Ex11 and Ex12 is essential for RNA editing of the AZIN1 coding site, and In11 prevents this RNA editing by inhibiting the formation of a stable dsRNA structure.

**Figure 4.**
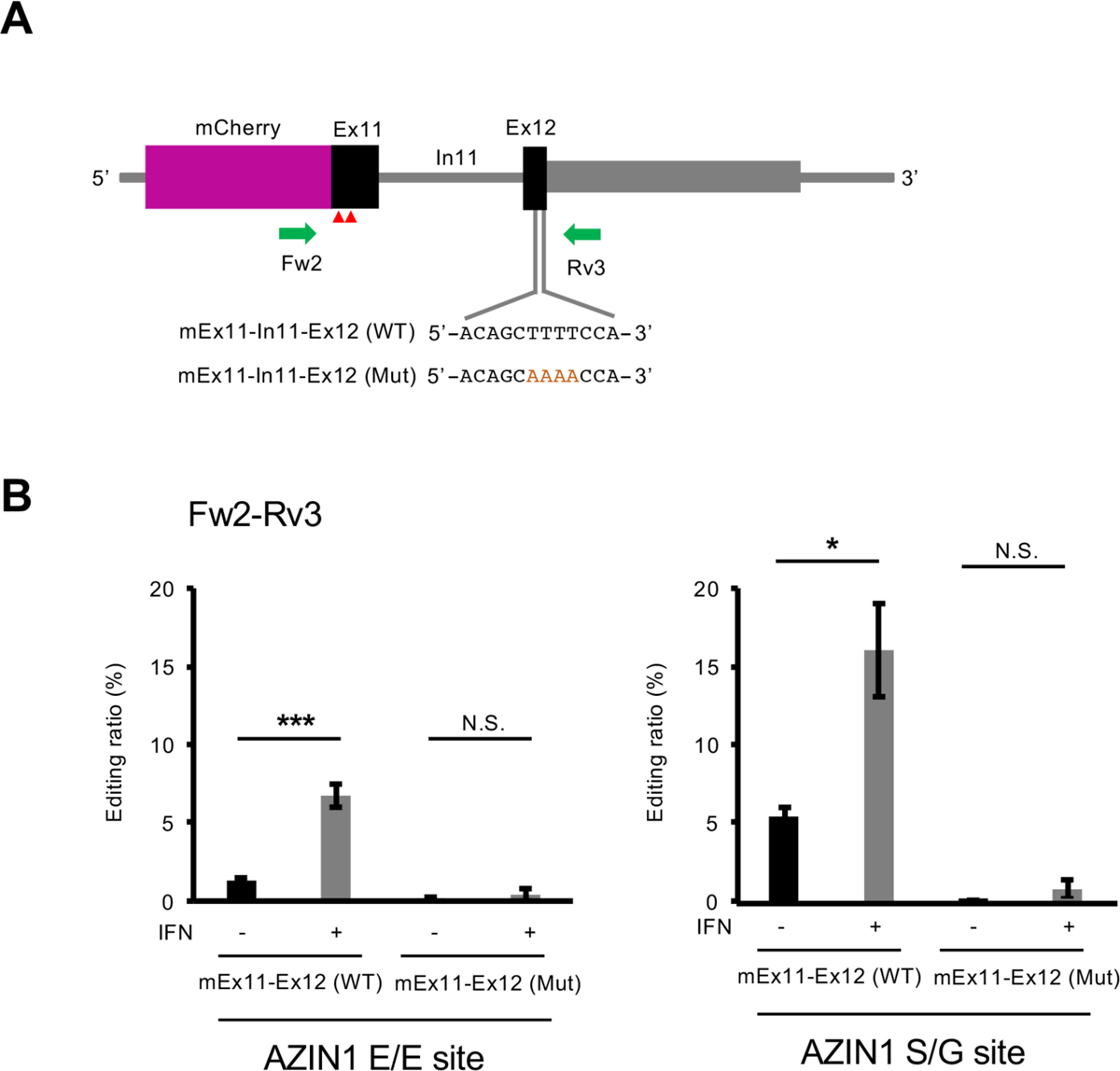
Identification of an editing complementary sequence in a downstream exon. (**A**) Schematic diagram of a reporter construct of the mouse *Azin1* gene encompassing exon 11 (Ex11), intron 11 (In10), and partial Ex12 containing only the coding region, which is fused to the *mCherry* gene. The position of PCR primers, and the position of point mutations (red AAAA) inserted in the editing complementary sequence in Ex12 are shown below. Two editing sites are indicated with red arrowheads. (**B**) After either an mCherry-fused mEx11-In11-Ex12 (WT) reporter vector or mEx11-In11-Ex12 (Mut) reporter vector, which contains four point mutations in Ex12, was transfected into Raw 264.7 cells, cells were treated with interferon (IFN) β1. Then, RNA was extracted and subjected to RT followed by PCR using Fw2-Rv3 primer sets. The PCR products derived from mature Azin1 reporter mRNA were used for the quantification of editing ratios at AZIN1 E/E and S/G sites in the absence or presence of IFNβ1. Values represent the mean ± SEM (n = 3 for each group; Student’s *t*-test, \**p* < 0.05, \*\*\**p* < 0.005, N.S., not significant).

### Intracellular localization of ADAR1 affects Azin1 RNA editing

ADAR1 p150 possesses a Zα domain and a NES in the N-terminus, which is not present in ADAR1 p110 (**Fig. 5A**). Given that Zα domain–mediated Z-RNA binding affects the RNA editing activity of ADAR1 p150 *in vitro* and *in vivo* (18,48), we first examined whether the Z-RNA binding capacity of ADAR1 p150 affects Azin1 RNA editing. For this purpose, we introduced tryptophan-to-alanine substitution at the position of amino acid 197 (W197A) in the Zα domain, which is known to lose binding ability to Z-formed nucleic acids (18,49), and expressed an EGFP-tagged mADAR1 p150 W197A mutant in A1/A2 dKO Raw 264.7 cells. As we previously reported (18), the W197A mutation did not affect the cytoplasm-dominant localization of ADAR1 p150 (**Fig. 5B**). After sorting GFP-positive cells by adjusting the expression level of wild-type ADAR1 p150 and W197A mutant proteins using the intensity of GFP fluorescence, we compared RNA editing levels. This analysis revealed that no significant alteration of RNA editing at two sites of Azin1 mRNA was detected (**Fig. 5C**). This finding indicates that Z-RNA binding capacity is dispensable for Azin1 RNA editing.

**Figure 5.**
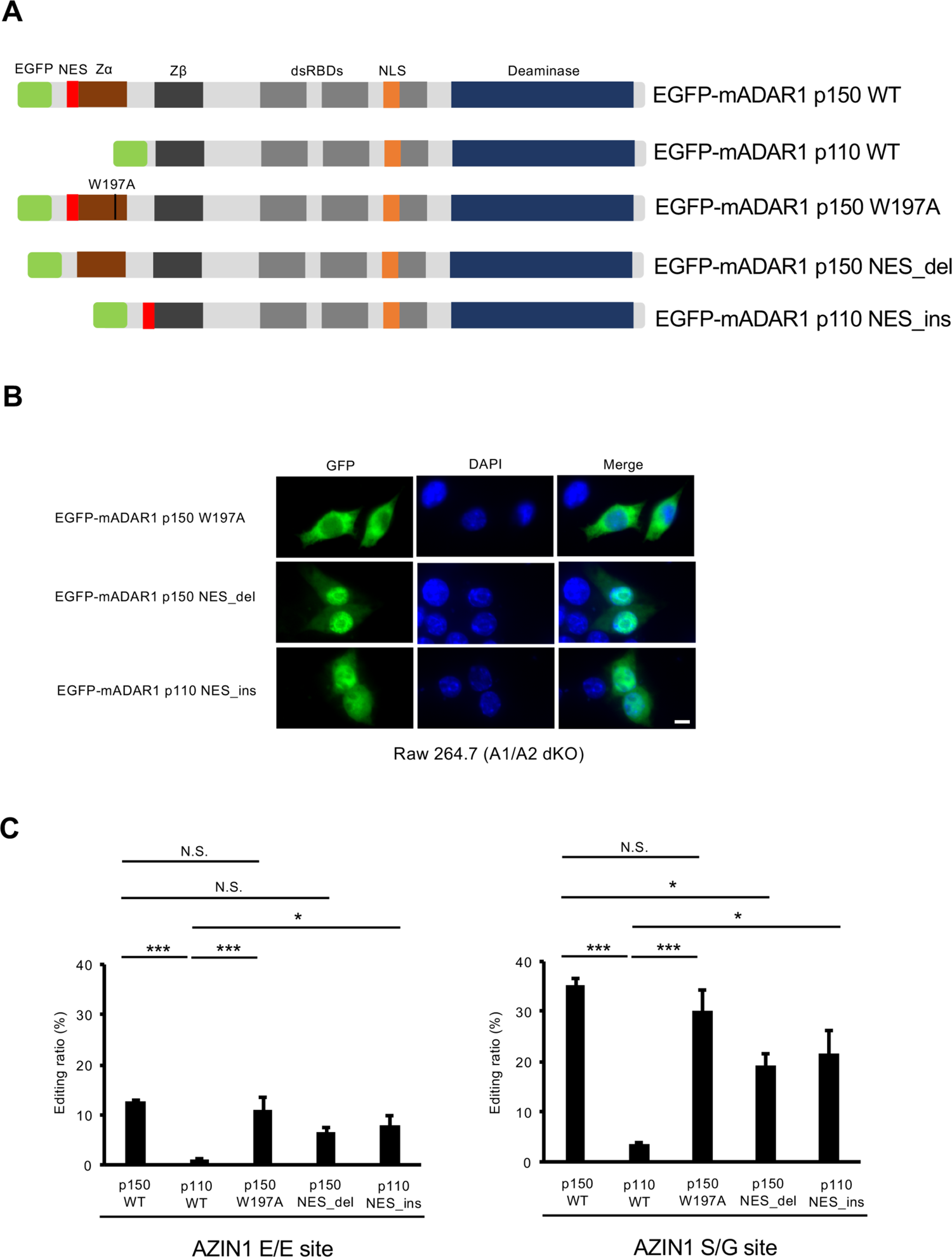
Intracellular localization of ADAR1 affects RNA editing of Azin1 mRNA. (**A**) Schematic diagram of EGFP-fused wild-type (WT) isoforms of mouse ADAR1 (mADAR1 p150 and p110) and their mutant isoforms. ADAR1 p150 and p110 share a Z-DNA/RNA-binding domain β (Zβ), three double-stranded RNA (dsRNA)-binding domains (dsRBDs), a deaminase domain (Deaminase) and a nuclear localization signal (NLS). In contrast, the Zα domain and a nuclear export signal (NES) are present only in the N-terminus of ADAR1 p150. Tryptophan at the position of amino acid 197 is substituted to alanine in the Zα domain of an EGFP-mADAR1 p150 W197A mutant, which leads to a loss of binding capacity to Z-DNA/RNA. The NES is deleted in an EGFP-mADAR1 p150 NES_del mutant, while the NES is inserted into the N-terminus of an EGFP-mADAR1 p110 NES_ins mutant. (**B**) The intracellular localization of indicated EGFP-fused mADAR1 mutants in *Adar1*/*Adar2* double-knockout (A1/A2 dKO) Raw 264.7 cells was visualized by GFP fluorescence. Nuclei were stained with DAPI. Merged fluorescent images are shown in the right panels. Scale bar, 10 μm. (**C**) After the indicated EGFP-tagged ADAR wild-type and mutant isoforms were expressed in A1/A2 dKO Raw 264.7 cells, RNA was extracted and subjected to reverse transcription (RT) followed by PCR using Fw1-Rv1 primer sets shown in Figure 1A. The PCR products derived from mature Azin1 mRNA were used for the quantification of editing ratios at two sites of Azin1 mRNA. Values represent the mean ± SEM (n = 3 for each group; Tukey’s honest significant difference test, \**p* < 0.05, \*\*\**p* < 0.005, N.S., not significant).

The intracellular localization of mADAR1 p150 is affected by multiple domains, including a NES, a part of a Zα domain, a NLS, and a part of the C-terminus (9,45,50,51). Thus, deleting a NES can partially but not completely shift ADAR1 p150 from the cytoplasm to the nucleus (9). As expected, we observed a partial shift of EGFP-tagged mADAR1 p150 NES_del mutant, in which a NES was deleted, to the nucleus (**Fig. 5A, B**). Then, we quantified RNA editing levels after adjusting the expression level of wild-type ADAR1 p150 and NES_del mutant using the intensity of GFP fluorescence. This demonstrated that RNA editing levels at two coding sites were significantly reduced (**Fig. 5C**). Next, we inserted a NES to the N-terminus of ADAR1 p110 (**Fig. 5A**) and expressed an EGFP-tagged mADAR1 p110 NES_ins mutant in A1/A2 dKO Raw 264.7 cells. A part of the mADAR1 p110 NES_ins mutant was observed in the cytoplasm and not detected in the nucleolus where wild-type ADAR1 p110 is localized (**Fig. 2A and 5B**). Accordingly, we found that RNA editing levels at two coding sites were significantly increased (**Fig. 5C**). Taken together, the intracellular localization of ADAR1, but not binding capacity to Z-RNA, is critical for regulating Azin1 RNA editing.

### ADAR1 p150 is an enzyme responsible for Azin1 RNA editing *in vivo*

We recently reported that editing levels at two sites of Azin1 mRNA are not reduced in *Adar1 p110^-/-^/Adar2^-/-^* mice (14), which suggests that ADAR1 150 is responsible for Azin1 RNA editing *in vivo*. To reciprocally reconfirm this finding, we generated *Adar1 p150*–specific KO (*Adar1 p150^-/-^*) mice by deleting the type I IFN-inducible promoter and exon 1A, as previously reported (21), using a genome editing system (**Fig. 6A**). We found that *Adar1 p150^-/-^* mice showed embryonic lethality, which was rescued by concurrent deletion of the *Ifih1* gene as previously observed (21,24). We then extracted RNA from the brain, thymus, and spleen at P0 and further confirmed that no Adar1 p150 mRNA was detected, whereas the expression level of Adar1 p110 mRNA was sustained in *Adar1 p150^-/-^ Ifih1^-/-^* mice (**Fig. 6B, C**). Accordingly, we observed that ADAR1 p150 proteins were selectively lost in all organs of mutant mice examined at P0 (**Fig. 6D**). Given that RNA editing of Blcap coding sites requires ECS in the upstream intron (6,14,34), the *in vitro* editing assay revealed that these sites are editable by nuclear ADAR1 p110 and ADAR2 but not ADAR1 p150 before splicing (**Fig. S1C**). Accordingly, although Blcap RNA editing was absent in *Adar1 p110^-/-^/Adar2^-/-^* mice (14), the editing levels of the two sites in Blcap mRNA in multiple organs of the *Adar1 p150^-/-^ Ifih1^-/-^* mice examined were comparable to those in wild-type mice (**Fig. 6E, F**). In contrast, the editing level of Azin1 coding sites was substantially absent (less than 1%) in all organs of *Adar1 p150^-/-^ Ifih1^-/-^* mice examined at P0 (**Fig. 6G, H**). In addition, we found that E/E and S/G sites were edited by less than 6% in the brain of wild-type mice where ADAR1 p110 is abundant, whereas ADAR1 p150 was expressed at its lowest level (**Fig. 6D, G, H**) as we previously reported (14–17). Taken together, these data indicate that ADAR1 p150 mediates Azin1 RNA editing *in vivo* and *in vitro*, which cannot be compensated by ADAR1 p110 and ADAR2.

**Figure 6.**
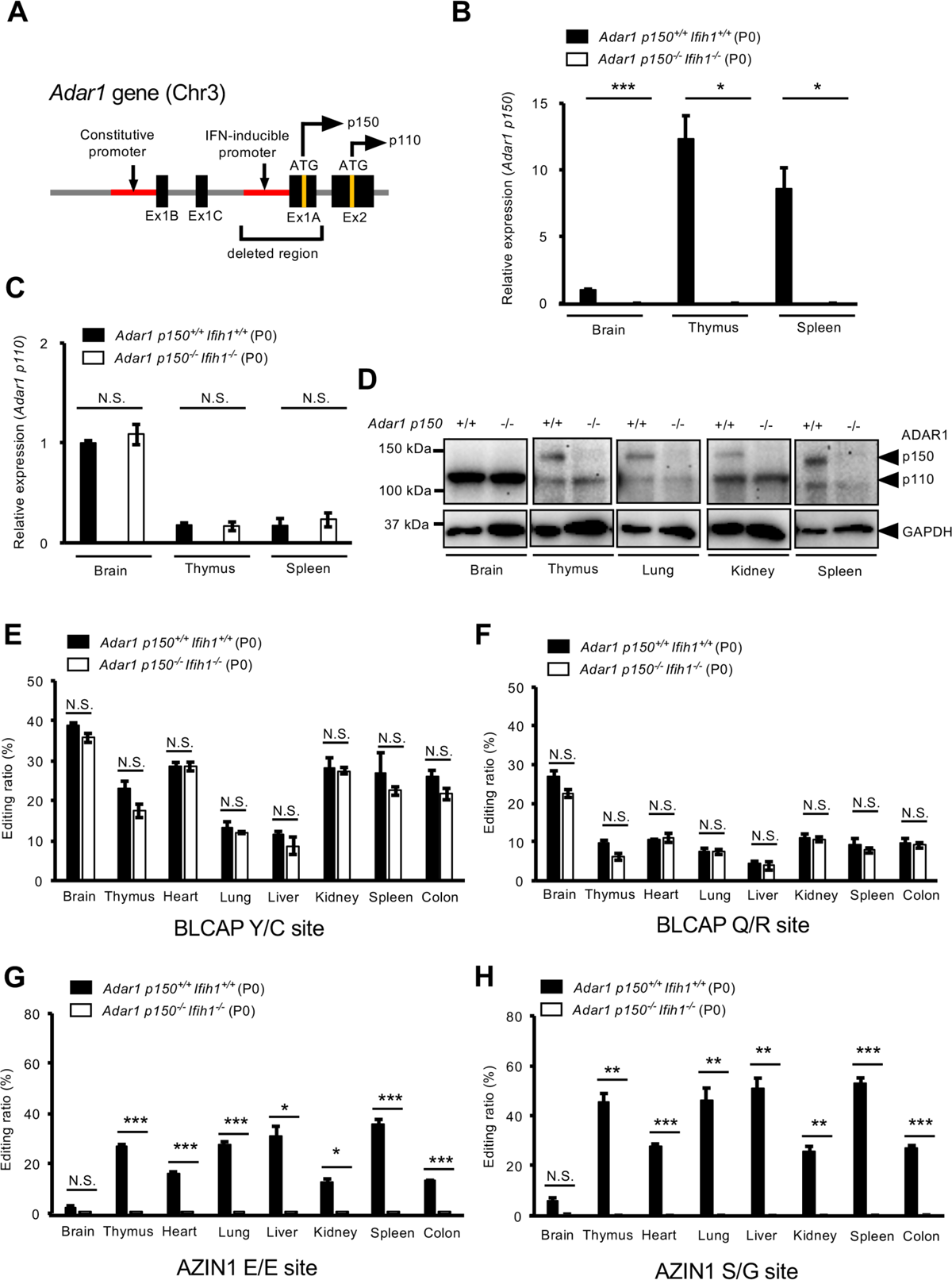
RNA editing of Azin1 mRNA is lost in *Adar1 p150*–specific KO mice. (**A**) Schematic diagram of the mouse *Adar1* gene. Adar1 p150 mRNA was transcribed from a type I interferon (IFN)-inducible promoter. Adar1 p110 transcripts usually contain exon 1B (Ex1B) or minor Ex1C and are translated from an initial methionine site (ATG) located in Ex2, while p150 transcripts contain Ex1A, which includes the initial methionine site. The genomic region deleted in *Adar1 p150*–specific knockout (*Adar1 p150^-/-^*, KO) mice is indicated. **(B, C)** The relative expression of Adar1 p150 mRNA (**B**) and Adar1 p110 mRNA (**C**) in the brain, thymus, and spleen at post-natal day (P0) was compared between *Adar1 p150^+/+^ Ifih1^+/+^* (wild-type) and *Adar1 p150^-/-^ Ifih1^-/-^* mice. Values represent relative gene expression normalized to GAPDH mRNA and are displayed as the mean ± SEM (n = 3 mice for each group; Student’s *t*-test, \**p* < 0.05, \*\*\**p* < 0.005, N.S., not significant). The mean values of Adar1 p150 (**B**) and Adar1 p110 (**C**) expression in the brains of wild-type mice were set as 1. (**D**) The expression of ADAR1 p110, and p150 proteins in various organs at P0 was compared between *Adar1 p150^+/+^ Ifih1^+/+^* and *Adar1 p150^-/-^ Ifih1^-/-^* mice using western blots. The expression of GAPDH protein is shown as a reference. (**E–H**) Editing ratios at BLCAP Y/C (**E**), BLCAP Q/R (**F**), AZIN1 E/E (**G**), and AZIN1 S/G (**H**) sites at P0 in various organs were compared between *Adar1 p150^+/+^ Ifih1^+/+^* and *Adar1 p150^-/-^ Ifih1^-/-^* mice. Values are displayed as the mean ± SEM (n = 3**–**6 mice for each group; Student’s *t*-test, \**p* < 0.05, \*\**p* < 0.01, \*\*\**p* < 0.005, N.S., not significant).

## Discussion

AZIN1 was originally identified from rat liver extract as a regulator of the intracellular level of polyamines (52,53). AZIN1 has two conserved RNA editing sites in the coding region (synonymous E/E and non-synonymous S/G sites), which were originally identified in human transcripts (33,54). Serine (S)-to-glycine (G) substitution at this site is predicted to affect protein conformation, inducing a cytoplasmic-to-nuclear translocation (55). Although knowledge on the physiological significance of such RNA editing is limited, it was recently reported that the S/G site was highly edited in hematopoietic stem and progenitor cells, which promotes the differentiation of these cells by altering the chromatin distribution of DEAD box polypeptide 1 in the nucleus (56). Of note, the maintenance of hematopoietic stem cells requires RNA editing mediated by ADAR1, especially ADAR1 p150, although protein recoding by ADAR1-mediated RNA editing is dispensable for normal development (18,22,57,58). Therefore, further investigation is needed to elucidate the role of RNA editing at the S/G site of AZIN1 in hematopoiesis *in vivo*.

We previously reported that RNA editing of Azin1 was absent in *Adar1^E861A/E861A^* mice, whereas the editing level is sustained in *Adar2^-/-^* mice, indicating that ADAR1 is responsible for RNA editing of Azin1 *in vivo* (6). Given that these coding sites are generally edited in the nucleus, one group reported that nuclear ADAR1 p110 was the responsible enzyme (55,59). Furthermore, the same group recently reported that AZIN1 RNA editing required a dsRNA structure formed between Ex11 and Ex12 in the presence of In11 in the nucleus (60). In contrast, we found that editing levels at two sites of Azin1 mRNA were not reduced in *Adar1 p110^-/-^/Adar2^-/-^* mice (14), which suggests that ADAR1 150 is responsible for Azin1 RNA editing *in vivo*. In addition, Kleinova et al recently reported that AZIN1 coding sites were edited by ADAR1 p150 but not ADAR1 p110 *in vitro* (61). However, the molecular mechanisms underlying ADAR1 p150–specific RNA editing of AZIN1 coding sites remain unresolved. We demonstrated for the first time that AZIN1 RNA editing is mediated by ADAR1 p150 after the splicing of In11, which acts as a potent suppressor for RNA editing. In addition to the short dsRNA (∼50 bp) formed between Ex11 and Ex12, In11 has sequences that might hybridize with Ex11 (55), which likely inhibits the formation of a stable dsRNA structure in the nucleus. Furthermore, in some cell lines, such as HEK293 and PLC8024, the editing level at AZIN1 coding sites is negligible, although total ADAR1 expression is high (55,62). This discrepancy is attributable to the relatively higher expression of ADAR1 p110 in these cells, at least in part, since we also observed the same pattern in HEK293T cells. Collectively, we propose that AZIN1 coding sites, which are conserved between humans and mice, can be used as distinct biomarkers for the RNA editing activity of ADAR1 p150.

Many studies suggest that RNA editing at the S/G site of AZIN1 is increased in multiple cancers, such as hepatocellular carcinoma, esophageal squamous cell carcinoma, gastric cancer, and colorectal cancer (55,59,63,64). Edited AZIN1, which is translocated into the nucleus, promotes cell proliferation and tumour progression through polyamine-dependent and -independent mechanisms (52,55). In addition, edited AZIN1 promotes cancer invasion, migration, and stemness in colorectal cancers (63). The elevated RNA editing at the S/G site of AZIN1 is most likely a response to type I IFN production from the chronic inflammatory environment of cancers in addition to a possible increase in *ADAR1* copy numbers (65). Furthermore, certain cancer cells acquire the ability to produce type I IFN and these ISG signature–positive cancer cells are sensitive to ADAR1 depletion (66–68). Of note, ADAR1 p150 but not ADAR1 p110 is responsible for lethality in cancer cells induced by ADAR1 loss (66). These mechanisms are independent of MDA5-sensing pathways but partially depend upon the activation of protein kinase R (PKR), which suggests the presence of other pathways underlying lethality caused by a loss of ADAR1 p150. Therefore, reduced RNA editing at the S/G site of AZIN1 might contribute to this mechanism, and therefore this needs further investigation.

## Experimental procedures

### Ethics statement

All experimental procedures that included mice were performed by following protocols approved by the Institutional Animal Care and Use Committee of Osaka University (27-004-023 and 01-063-005).

### Mice maintenance

Mice were housed with food and water available ad libitum in a light (12 hours on/12 hours off), temperature (23 ± 1.5°C), and humidity (45 ± 15%) controlled environment as previously described (18,69).

### Mutant mice

*Ifih1^-/-^* mice were maintained in our laboratory as previously described (16,18). *Adar1 p150^-/-^* mice were generated by genome editing using a CRISPR/Cas9 system at the Genome Editing Research and Development Center, Graduated School of Medicine, Osaka University as previously described (14). Briefly, two Alt-R CRISPR-Cas9 CRISPR RNAs (crRNAs; CrRNA-Adar1p150-KO-up [5’-TTGTAAATCTCGCAAGCAGT-3’] and CrRNA-Adar1p150-KO-down [5’-GCTGGCAGTTCGGCTTGAGA-3’]) were synthesized at Integrated DNA Technologies (IDT) and hybridized with trans-activating CRISPR RNA (tracrRNA), generating guide RNAs, which were introduced into pronuclear-stage mouse embryos with Cas9 mRNA by electroporation. Mouse embryos that developed to the two-cell stage were transferred into the oviducts of female surrogates. Genotyping PCR of samples from *Adar1 p150^-/-^* mice was performed using the following three primers: 5’-AAACGCATCAGGTACCCAGG-3’, 5’-CTCCGCCCTGTGAGGAAGTT-3’, and 5’-CAGCTGGGGCTCATGTACGA-3’, which generated two fragments with different sizes: one for a wild-type allele and another for a mutated allele. All mice used in experiments were in a C57BL/6J background.

### Cell culture

Mouse macrophage-derived Raw 264.7 and human HEK293T cell lines were cultured in Dulbecco’s modified Eagle’s medium (DMEM) (Nacalai Tesque) containing 10% (v/v) fetal bovine serum at 37°C in the presence of 5% CO_2_. A1/A2 dKO Raw 264.7 cells were established in a previous study (18).

### Construction of plasmids

The following plasmids, pEGFP-C1-mADAR1 p150, pEGFP-C1-mADAR1 p150 W197A, and pEGFP-C1-mADAR1 p110 were previously reported (18). Similarly, we generated pEGFP-C1-mADAR2 by inserting the coding region of mouse ADAR2, which was amplified from a plasmid containing the mouse *Adar2* gene obtained from DNAFORM (Yokohama, Japan), into a pEGFP-C1 vector using EcoRI/BamHI restriction enzyme recognition sites. To create pEGFP-C1-mADAR1 p150 NES_del and pEGFP-C1-mADAR1 p110 NES_ins plasmids, site-directed mutagenesis was performed by using a PrimeSTAR mutagenesis basal kit (Takara Bio) and pEGFP-C1-mADAR1 p150 as a template with the following primers: 5’-TGCTGACCAGAGTCCGGAGCAGAAG-3’ and 5’-GGACTCTGGTCAGCACCTCTCCATGG-3’, and 5’-CATCAGTGCTGAAATCAAGGAGAAG-3’ and 5’-ATTTCAGCACTGATGCTCAGCTCCCG-3’, respectively. The coding region of human ADAR1 p110, ADAR p150, and ADAR2 were amplified by PCR from plasmids containing these genes, which were obtained from Open Biosystems. The resultant PCR products were inserted into a pFN21A HaloTag CMV Flexi Vector (Promega) using SgfI/PmeI restriction enzyme recognition sites, which were termed pHaloTag-hADAR1 p150, pHaloTag-hADAR1 p110, and pHaloTag-hADAR2, respectively. The corresponding regions of a partial mouse *Azin1* gene were amplified from mouse tail genomic DNA by using Phusion Hot Start Flex 2X Master Mix (New England Biolabs) and the following primers: 5’-TATTCTCGAGTAGAAAAAAATGGGAGTGAT-3’ and 5’-AATAATGGATCCCAATCACTGAATGACATCAT-3’ for mEx10-In10-Ex11, 5’-TATTCTCGAGCCAAATACAAGGAAGATGAGCC-3’ and 5’-AATAATGGATCCCAATCACTGAATGACATCAT-3’ for mEx11, and 5’-TATTCTCGAGCCAAATACAAGGAAGATGAGCC-3’ and 5’-AATAATGGATCCAACCCAGTTAATGGGCTTCCA-3’ for mEx11-In11-Ex12. The resultant PCR products were inserted into a pmCherry-C1 (Clontech) using XhoI/BamHI restriction enzyme recognition sites. To insert four point mutations (underlined in the primer sequences) in Ex12 of a mEx11-In11-Ex12 reporter construct, site-directed mutagenesis was performed using a PrimeSTAR mutagenesis basal kit (Takara Bio) with the following primers: 5’-AGACAGC<u>AAAA</u>CCACTGAAGCTTAAACAG-3’ and 5’-TCAGTGG<u>TTTT</u>GCTGTCTTCTTGGCTCAG-3’. All the constructs were verified by Sanger sequencing.

### Plasmid transfection

The following plasmids, pEGFP-C1-mADAR1 p150, pEGFP-C1-mADAR1 p150 W197A, pEGFP-C1-mADAR1 p150 NES_del, pEGFP-C1-mADAR1 p110, pEGFP-C1-mADAR1 p110 NES_ins or pEGFP-C1-mADAR2, were transfected into A1/A2 dKO Raw 264.7 cells using a Neon Transfection System (Thermo Fisher Scientific) with the following parameters: pulse voltage, 1,750 V; pulse width, 25 ms; and pulse number, 1, as previously described (18). Twenty-four hours after transfection, GFP-positive cells were sorted by adjusting the expression of each mADAR protein using the intensity of GFP fluorescence and an SH800 cell sorter (Sony) for subsequent RNA extraction. For microscopic analyses, A1/A2 dKO Raw 264.7 cells were maintained in chamber slides and plasmids were transfected using Lipofectamine 2000 (Thermo Fisher Scientific) in presence of NATE^TM^ (InvivoGen), a transfection enhancer. Twenty-four hours after transfection, cells were fixed with 4% paraformaldehyde for 30 min at room temperature and stained with DAPI Fluoromount-G (Southern Biotech) to observe intracellular localization using an Olympus BX63 fluorescence microscope. For treatment with type I IFN, Raw 264.7 cells were cultured with 100 ng/mL recombinant mouse IFNβ1 (BioLegend). After 24 h, partial mouse *Azin1* expression plasmids were transfected using Lipofectamine 2000 in the presence of NATE^TM^ (InvivoGen) in some experiments.

### Preparation of tissue and cell lysates

Tissue lysates were prepared as previously described (18). In brief, isolated organs were frozen in liquid nitrogen, thawed once at room temperature, and then homogenized in lysis buffer 1 (0.175 M Tris-HCl, pH 6.8, 3% SDS, and 5 mM EDTA). After boiling at 95°C for 10 min, the lysates were subjected to centrifugation at 20,000 × g and 4°C for 10 min. Each supernatant was transferred to a 1.5 mL tube and stored at −80°C until use. To prepare cell lysates, collected cells were lysed with lysis buffer 2 (20 mM Tris-HCl, pH 7.9, 25% glycerol, 420 mM NaCl, 1.5 mM MgCl_2_, 0.2 mM EDTA, 0.5 mM PMSF, 0.5 mM DTT), frozen in liquid nitrogen and then thawed on ice five times. After centrifugation at 20,000 × g and 4°C for 10 min, the supernatants were transferred to new 1.5-mL tubes and stored at −80°C until use.

### Immunoblot analysis

Each lysate was separated by sodium dodecyl sulfate–polyacrylamide gel electrophoresis and transferred to polyvinylidene difluoride membranes (Bio-Rad). Immunoblotting was performed by using the following primary antibodies: mouse monoclonal anti-ADAR1 antibody (15.8.6; Santa Cruz Biotechnology), mouse monoclonal anti-ADAR2 antibody (1.3.1; Santa Cruz Biotechnology), rabbit polyclonal anti-AZIN1 antibody (11548-1-AP; Proteintech), mouse monoclonal anti-GFP antibody (GF200; Santa Cruz Biotechnology), rabbit polyclonal anti-Halo Tag antibody (G928A; Promega), and mouse monoclonal anti-GAPDH antibody (M171-3; MBL). Chemiluminescence was detected by using ImmunoStar Zeta (Fujifilm) or Pierce ECL Western Blotting Substrate (Thermo Fisher Scientific).

### RNA extraction

Total RNA was extracted from isolated organs or collected cells using TRIzol reagent (Thermo Fisher Scientific) by following the manufacturer’s instructions. After measuring the RNA concentration using a NanoDrop One (Thermo Fisher Scientific), total RNA samples were stored at −80°C until use.

### Quantification of the RNA editing ratio with Ion amplicon sequencing reads

The preparation of Ion amplicon libraries for the quantification of RNA editing sites has been previously described (6,14). In brief, 500 ng of total RNA was incubated with 0.1 U/µL DNase I (Thermo Fisher Scientific) at 37°C for 15 min and then denatured at 65°C for 15 min. The resultant RNA samples were reverse transcribed into cDNA using a SuperScript III First-Strand Synthesis System (Thermo Fisher Scientific) with oligo (dT). Synthesized cDNA was then amplified using Phusion Hot Start Flex 2X Master Mix (New England Biolabs) and the following primers: human endogenous AZIN1, 5’-TCGCAGTTAATATCATAGC-3’ and 5’-GCTTCAGCGGAAAAGCTGTC-3’; mouse endogenous Azin1, 5’-GATGAGCCAGCCTTCGTGT-3’ (Fw1) and 5’-TGGTTCGTGGAAAGAATCTGC-3’ (Rv1) or 5’-ACCAGCAAATCTAAACTGTCACT-3’ (Rv2); mCherry-fused partial AZIN1 expression construct, 5’-CACTACGACGCTGAGGTCAA-3’ (Fw2) and 5’-GGGAGGTGTGGGAGGTTTT-3’ (Rv3) or 5’-ACCAGCAAATCTAAACTGTCACT-3’ (Rv4). A second round of PCR was then performed using an aliquot of the first PCR product as a template and second primers that were editing-site specific; an A adaptor (5’-CCATCTCATCCCTGCGTGTCTCCGACTCAG-3’), an Ion Xpress Barcode™ and a trP1 adaptor (5’-CCTCTCTATGGGCAGTCGGTGAT-3’) were in forward and reverse primers, respectively. The sequences of the second primers for human endogenous AZIN1 and mouse endogenous Azin1 were previously described (6,14). The primer sets for mouse Blcap and Nova1 were previously described (6). After gel purification, the concentration of each PCR product was measured using a NanoDrop One and then equal amounts of approximately 200-bp PCR products were combined. After a quality examination using a 2100 Bioanalyzer (Agilent Technologies) with a High Sensitivity DNA kit, the resultant amplicon library samples were subjected to deep sequencing using an Ion S5 system (Thermo Fisher Scientific) at the CoMIT Omics Center, Graduate School of Medicine, Osaka University. RNA editing ratios were calculated using an in-house program as previously described (6,14).

### qRT–PCR

As previously described (16,18,69), cDNA was synthesized from total RNA extracted from various organs and cells. In brief, after 500 ng of each total RNA sample was denatured at 65°C for 5 min, cDNA was synthesized using a ReverTra Ace qPCR–RT Master Mix with guide DNA Remover (Toyobo). The quantitative reverse transcription (qRT)–PCR reaction mixture was prepared by combining each target-specific primer and probe with a THUNDERBIRD Probe qPCR Mix (Toyobo). The qRT–PCR was performed using a QuantStudio™ 7 Flex Real-Time PCR System (Thermo Fisher Scientific). The sequences of primers and probes for Adar1 p150 and GAPDH have been previously reported (16), while the sequences for primers and probes of Adar1 p110 and Azin1 were obtained from IDT: Adar1 p110-primer1 (5’-GCTGAAGCTGGAAACTCCTA-3’), Adar1 p110-primer2 (5’-GCAGCGTCCGAGGAATC-3’), Adar1 p110-probe (5’-/56-FAM/AGTACGACT/ZEN/GTGTCTGGTGAGGGA/3IABkFQ/-3’); Azin1-primer1 (5’-TTCATCTCAGCCGTATTCCAC-3’), Azin1-primer2 (5’-GTTCCATCTCCTAACTTGTCCA-3’), Azin1-probe (5’-/56-FAM/TGGCCCGTC/ZEN/TCTTGTTTTTCCTGG/3IABkFQ/-3’). The expression level of each Adar and Azin1 mRNA relative to that of GAPDH mRNA were calculated by the ΔΔCt method.

### Analysis of dsRNA structure

Potential secondary dsRNA structures were calculated using a RNAfold web server (14,18,70).

### Statistical analyses

For statistical analyses, an unpaired two-tailed Student’s *t*-test or Tukey’s honest significant difference (HSD) test was used as indicated in each figure legend. All values are displayed as the mean ± standard error of the mean (SEM). No significance is displayed as N.S., while statistical significance is displayed as *p* < 0.05 (*), *p* < 0.01 (**) or *p* < 0.005 (***).

### Supporting information

This article contains supporting information.

## Acknowledgements

We thank Ms. Aki Irie, Ms. Akane Tejima and all staff at the Genome Editing Research and Development Center, the Center for Medical Research and Education, the CoMIT Omics Center, and the Institute of Experimental Animal Sciences, Graduate School of Medicine, Osaka University, for technical support. Computations were partially performed on an NIG supercomputer at ROIS National Institute of Genetics, Japan.

## Funding and additional information

This work was supported by Grants-in-Aid KAKENHI [20H03341 and 23H02576 to Y. Kawahara, 18K15186 and 21K07080 to T.N., 20J11266 to J.I.K., and 18K11526 to Y. Kato] from the Ministry of Education, Culture, Sports, Science and Technology (MEXT) of Japan, a grant [JP20ek0109433, JP21ek0109433 and JP22ek0109433 to T.N.] from the Japan Agency for Medical Research and Development (AMED), grants from the Japan Intractable Diseases Research Foundation, Astellas Foundation for Research on Metabolic Disorders, The Waksman Foundation of Japan, and a Kishimoto Fund research grant from Senri Life Science Foundation [to T.N.], and grants from The Takeda Science Foundation and the Uehara Memorial Foundation [to Y. Kawahara and T.N.]. This research was also supported by The Nippon Foundation–Osaka University Project for Infectious Disease Prevention [to Y. Kawahara]. F.X. was supported by MEXT scholarships. J.I.K. was supported by The Korean Scholarship Foundation and a Research Fellowship for Young Scientists from the Japan Society for the Promotion of Science (JSPS).

## Abbreviations

The abbreviations used are: ADAR, adenosine deaminase acting on RNA; AZIN1, antizyme inhibitor 1; dsRNA, double-stranded RNA; ECS, editing (or exon) complementary sequence; IFN, interferon; ISGs, interferon-stimulated genes; KI, knock-in; KO, knockout; MAVS, mitochondrial antiviral signaling protein; MDA5, melanoma differentiation-associated protein 5; NES, nuclear export signal; NLS, nuclear localization signal; UTR, untranslated regions; Zα, Z-DNA/RNA-binding domain α.

