## Supplemental Figures for "RNA editing of AZIN1 coding sites is catalyzed by ADAR1 p150 after splicing"

This file includes:

Supplementary Figures S1 to S3

**Supplementary Figures**

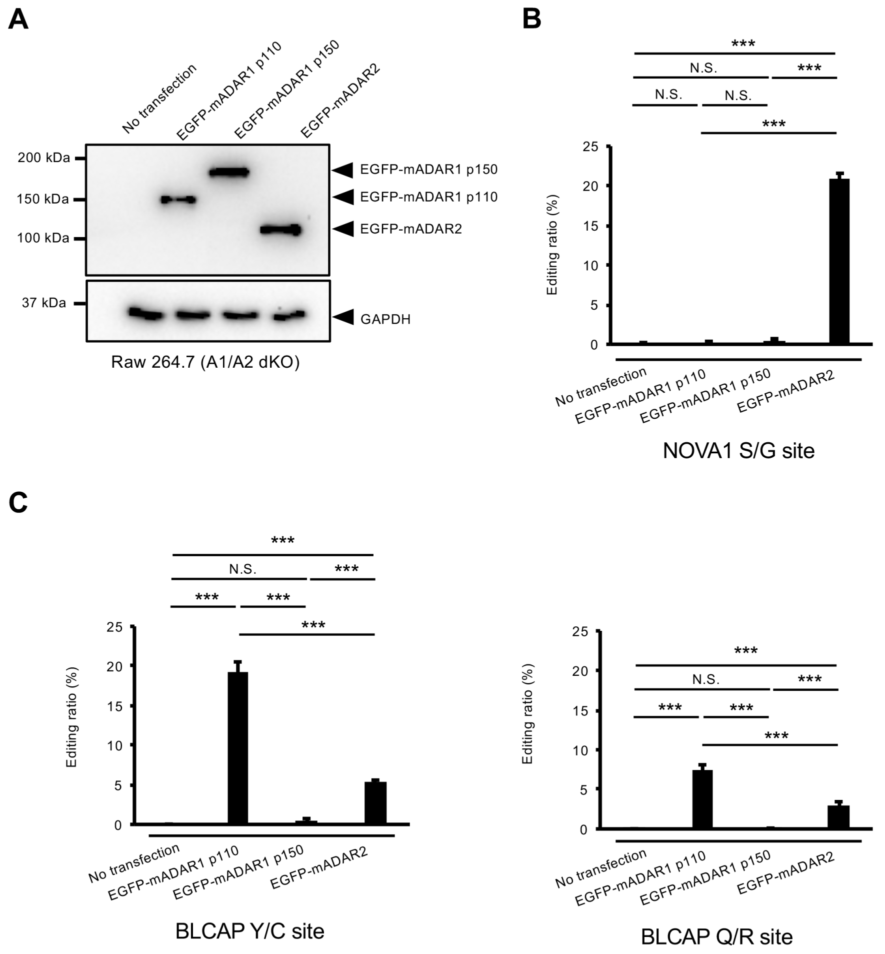


**Supplementary Figure S1.** ADAR isoform–specific RNA editing. (**A**) The expression of EGFP-tagged mouse ADAR1 p110 (mADAR1 p110), mADAR1 p150, and mADAR2 proteins in *Adar1*/*Adar2* double-knockout (A1/A2 dKO) Raw 264.7 cells was detected using anti-GFP antibody. The expression of GAPDH protein is shown as a reference. (**B, C**) Editing ratios at NOVA1 S/G (**B**), BLCAP Y/C, and BLCAP Q/R (**C**) sites were compared among indicated EGFP-tagged ADAR isoforms expressed in A1/A2 double knockout (dKO) Raw 264.7 cells. Values represent the mean ± SEM (n = 3 for each group; Tukey's honest significant difference test, ****p* < 0.005, N.S., not significant).

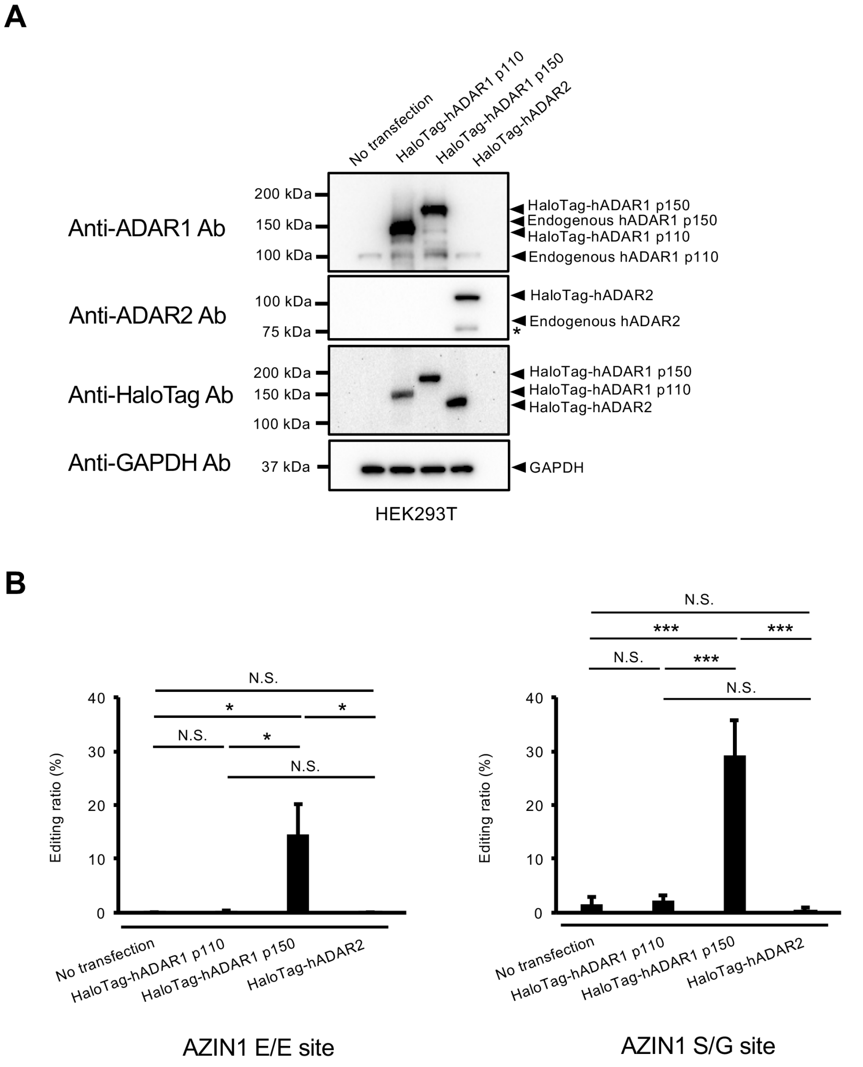


**Supplementary Figure S2.** ADAR1 p150–specific RNA editing of human AZIN1 mRNA. (**A**) The expression of HaloTag-fused human ADAR (hADAR) proteins expressed in human HEK293T cells was detected using anti-ADAR1, anti-ADAR2, or anti-HaloTag antibodies. The expression of GAPDH protein is shown as a reference. The truncated ADAR2 expression is indicated with an asterisk (*). (**B**) Editing ratios at AZIN1 E/E and S/G sites were compared among indicated HaloTag-fused hADAR isoforms expressed in HEK293T cells. Values represent the mean ± SEM (n = 3 for each group; Tukey's honest significant difference test, *$p < 0.05$, ***$p < 0.005$, N.S., not significant).

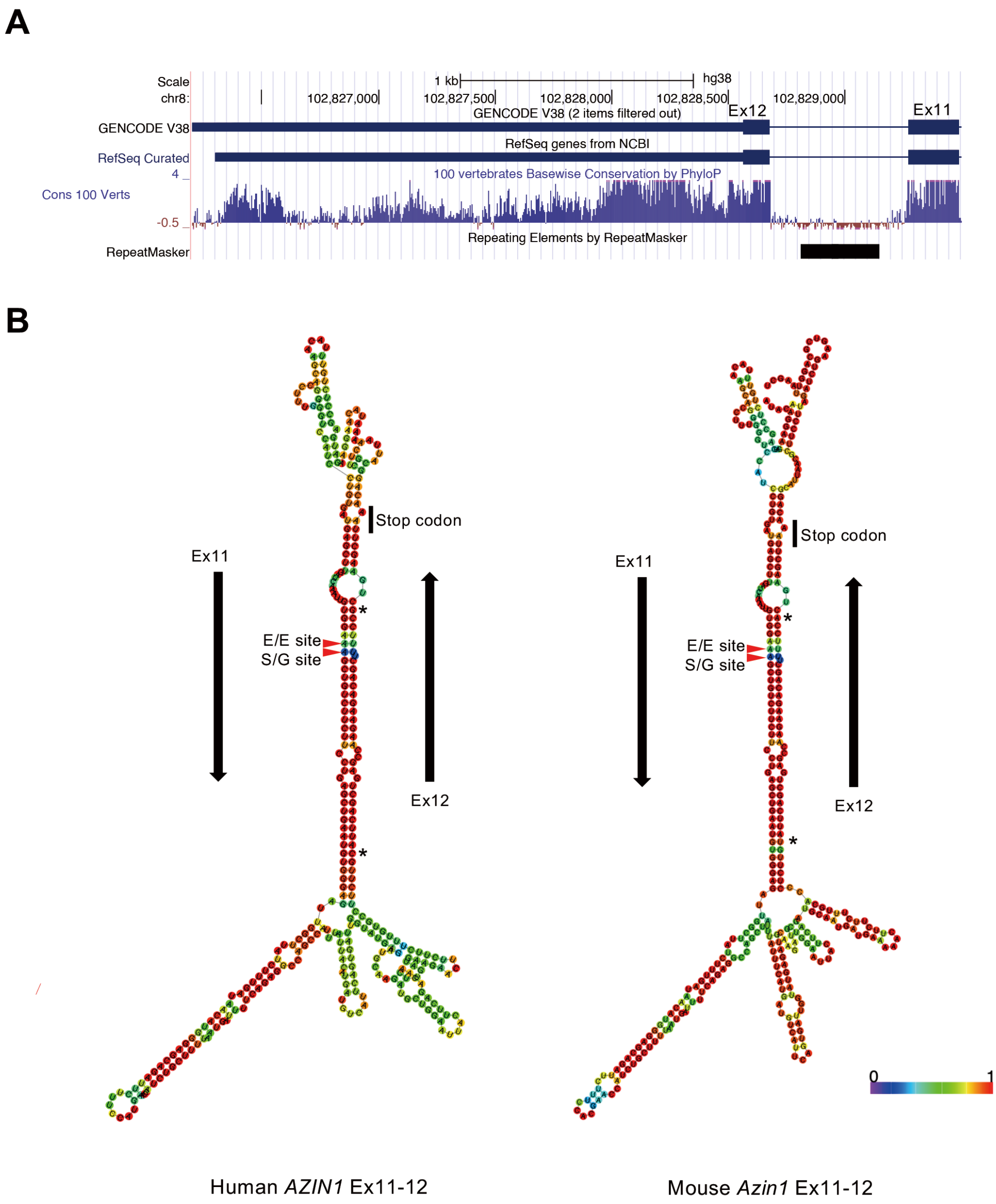


**Supplementary Figure S3.** Conservation of the secondary structure required for AZIN1 RNA editing between human and mouse. (**A**) The genomic region encompassing exon 11 (Ex11), intron 11, and exon 12 (Ex12) of human AZIN1 was analyzed using a University of California Santa Cruz (UCSC) genome browser. Conservation among 100 vertebrates is shown. (**B**) The secondary structure formed between Ex11 and partial Ex12 of human AZIN1 (left panel) or mouse Azin1 (right panel) mRNAs was estimated using a RNAfold web server and colored by base-pairing probabilities. The two editing sites are indicated by red arrowheads. Nucleotides that differ between human and mouse are indicated with an asterisk (*).
